# Metabolic profiling using benchtop NMR identifies the metabolomic signature of persistent CAR-T cells

**DOI:** 10.1101/2024.07.22.604702

**Authors:** Suk-Heung Song, Herb Ryan, Jens Hoefflin, Taeyoon Kyung, Joshua Mace, Jai Raman, Shawdee Eshghi

**Affiliations:** Ginkgo Bioworks, Boston, MA, USA

**Keywords:** metabolites, PCA, CARs, CAR-T cells, benchtop NMR

## Abstract

Analytical technologies for engineered biological systems hold great promise in addressing various challenges in modern pharmaceuticals and biomedical therapies. These endeavors often follow a design-build-test-learn approach, utilizing biological data from genetic circuits, signal pathways, metabolites, and proteins to optimize biological systems. Deciphering large and intricate datasets can prove to be a formidable task. Principal component analysis (PCA) tool is an invaluable method for reducing dataset complexity and enhancing interpretability while minimizing information loss, simultaneously. PCA tool achieves this by creating new, uncorrelated variables that capture the maximum variance in the data. Herein chimeric antigen receptors (CARs) T cells metabolomic study presents a slightly inverse problem, where PCA is applied to model sparse extracellular metabolites data from CAR-T cells, resulting in a two-component model. Using an benchtop NMR spectrometer only six metabolites could be annotated, nevertheless, the PCA model could identify differences in the metabolites of CAR-T cells based on the design of CARs, specifically the combinations of the intracellular domain (ICDs). It’s noteworthy that the behavior and fate of CAR-T cells are distinctly influenced by the type of ICDs used upon antigen recognition.

## 1. Introduction

Synthetic biology, SynBio has been accelerating in health & therapeutics, cultured ingredients, agriculture, flavors and fragrance, and biosecurity such as vaccines, because of reducing the cost of gene synthesis, and advances in genetic engineering, and data science. In their fields of study, large-scale data has become increasingly widespread. As the information on SynBio becomes more sophisticated, the size and complexity of data about their designs increase. To manage and interpret such datasets, methods are required that can significantly reduce their dimensionality while preserving as much information as possible. The optimal approach for this is to preserve as much statistical information as possible while reducing the dimensionality of a dataset. This means that the preservation of statistical information involves finding new variables. The new variables are linear functions of those in the original dataset. These linear functions can successively maximize variances that are uncorrelated from each other. Therefore, finding such new variables, such as principal components, can be accomplished by solving a linear algebra eigenvalue or vector problem.

Consequently, PCA tools are a statistical procedure that enables the summarization of information content in large data tables using a smaller set of summary indices, i.e., principal components, that are more easily visualized and analyzed. Datasets that can be analyzed using these tools can include compound or reagent reaction measurement data, point data from a continuous process, data from a batch process, biological entity data, or test data from a design of experiments protocol. The PCA is a multivariate statistical tool extensively used in the areas of data processing, pattern recognition, and signal processing, such as regression, classification, MS/LC, or NMR spectra, respectively. It is also a statistical method of factor analysis that forms the basis of multivariate data analysis based on projection methods. They can represent a multivariate data table as a smaller set of variables to observe trends, jumps, clusters, and outliers. The benefits of using them include the ability to uncover relationships between observations and variables, as well as among variables.

Historically, PCA tools have a rich history dating back to Pearson and Hotelling [1–2]. Since its inception, these tools have gained popularity and have been adapted in various fields. Pearson initially described the analysis as finding lines and planes that best-fit systems of points in space [3–4], and many publications have been dedicated to variants of them for specific types of data [5–6]. These tools are a statistical procedure that can be used to analyze datasets containing multicollinearity, missing values, categorical data, and imprecise measurements. PCA is aimed to extract information from a dataset and represent it as a set of summary indices called principal components [7]. It finds lines, planes, and hyperplanes in a K-dimensional space that approximates all data as closely as possible in the least squares sense. A line approximates the dataset point in the least squares sense, and then each point of the data maximizes the variance of the coordinates on the line. We begin by introducing the basic algorithm of PCA, apply this algorithm to analyze CAR-T cells’ metabolic 1H NMR spectral data, and characterize the differences in the metabolites of T cells according to CAR design. Recently this therapy is one of the promising cancer treatments that the FDA has approved and defined as engineering patients’ immune cells to treat their cancers. This is a new cancer therapy that uses engineered CAR-T cells as a part of the human immune system to target specific antigens [8–9]. Treatment has been aimed mainly at clinical responses, but many challenges limit the therapeutic efficacy. Several studies have investigated data sets that describe metabolites in the tumor microenvironment (TME) of CAR-T cells in clinical immunotherapy [10–12]. The high metabolic demands of tumor cells can compromise the effectiveness of CAR-T cells by competing with each other for nutrients in the TME. Thus, it is essential to prepare CAR-T cells to beat the metabolic barriers imposed by the TME. The metabolic analysis is a critical mechanism to contribute to the immunosuppressive TME. It is necessary to understand T cell function to identify ways to metabolically armor CAR-T cells and increase clinical efficacy [13–14]. In this regard, relative measurements of intracellular metabolomics for cells are useful in identifying candidate CAR-T cell signal pathway changes. Previous studies show a new role for CAR engineering to control T cell metabolism as a key determinant of T cell effector and memory responses [15–17]. The study reported the influence of signaling domains of coreceptors 4-1BB (BBz CAR) and CD28 (28z CAR) on the metabolic characteristics of human CAR T cells. The 4-1BB in the CAR architecture promoted the outgrowth of CD8+ central memory T cells that had significantly enhanced respiratory capacity, increased fatty acid oxidation, and enhanced mitochondrial biogenesis. In contrast, the CD28 domains in the CAR architecture yielded effector memory cells with a genetic signature consistent with enhanced glycolysis. These results provide that BBz CAR-T cells use fatty acids as the predominant energy source, whereas 28z CAR-T cells rely more on a glycolytic-based metabolism, which are characteristics of memory T cells and natural effector, respectively. These findings suggest, at least in part, a metabolic insight into the CAR-T cell behaviors and fates by expressing different signaling domains and informing the design of future CAR [18–20].

In this study, a major challenge in CAR design is ensuring better tumor control at a high tumor burden, while newly designed CAR-T cells proliferate significantly more than previous BBz or 28z CAR T-cells. Therefore, we employed three different CAR ICD designs, CAR2, CAR4, and CAR10. Especially CAR2 was encoding the CD28(28z CAR) design, whereas CAR4 and CAR10 were encoding the CD40 and 4-1BB (BBz CAR) design, respectively. From our results of PCA, we obtained meaningful insight into the distinct metabolic profile of CAR2, CAR4, and CAR10-T cells and quantitative differences over the different types of ICDs. These results can support the design types of CAR-T cells by metabolic insights

## 2. Methods

### 2.1. NMR sample preparation

CAR-T cells and A549 cell cultures were collected respectively. CAR-T cells are CAR2s, CAR4s, and CAR10s according to different CAR ICD designs. On the other hand, A549 cells are lung carcinoma epithelial cells that constitute a cell line, shortly lung tumor cells. All CAR samples are supernatant collected 72 hrs post A549 tumor cells co-culture and the supernatant is kept in the freezer (−80°C) until NMR data acquisition. A standard methanol-water chloroform extraction was used to extract metabolites from the cells. The samples were prepared using 500uL of sample mixed with 500uL of an NMR buffer containing a chemical shift reference and analytical standard (TSP-2, 2, 3, 3-d4 (D,98% Sodium-3-Trimethylsilyl-Propionate, Cambridge Isotope Laboratories, Inc.). The NMR buffer was prepared with Mono- and Dibasic Potassium Phosphate at 0.5M concentration and a pH of 7. 1M solutions of both Mono- and Dibasic Potassium Phosphate were prepared and a mixture of 61.5mL dibasic and 38.5mL monobasic potassium phosphate was prepared and the pH was measured at pH 7 as the Henderson–Hasselbalch equation evaluates to for a pK of 6.86 at 25C.

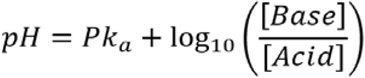

TSP-2, 2, 3, 3-d4 was added with a concentration of 30mM (516.81mG per 100mL added into a volumetric flask, and the pH 7 NMR buffer was added up to 100mL)

The samples were filled into standard Wilmad^®^ 5mM NMR tubes (MilliporeSigma, MA, USA) and an automated shimming routine was performed for each sample before measurement.

### 2.2 NMR spectra acquisition

CAR-T cell Spinsolve 80 MHz NMR (Magritek, Malvern, PA, USA) using 128 scans per sample, 15s relaxation delay, and water presaturation. All samples were acquired with an acquisition time of 6.4s and a saturation period of 2s. Free induction decays were multiplied by an exponential weighting function equivalent to 1 Hz line broadening after which they were Fourier transformed from the time to the frequency domain and referenced to the TSP single peak at 0.0 ppm. All spectra were then phased, and the baseline was corrected automatically. Areas of the spectrum <0 ppm and >10 ppm were excluded since they contained no data of interest. The spectra were converted into numerical vectors, representing the individual metabolites, by integrating across the spectrum using 0.04 ppm integral regions by using time-domain model-based quantitative NMR. The NMR spectral data were then converted to CSV format, normalized by sum, and applied concentration scaling. Individual metabolites were identified using standards, our library of NMR peak, and the human metabolome database (HMDB) 4.0 [21]. At this stage, the quantitative NMR process will be done with the multiplets sorted in the table based on their perceived ranking as candidates for compounds(i.e., target metabolites). The signal corresponding to the calibration compounds must be integrated and normalized according to the number of protons giving rise to the signal. The integrals of the compound of interest must then be compared with those of the calibration compound. The concentration of a compound x in the presence of a calibrant can be calculated using the following formula:

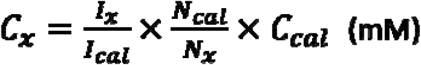

where *I, N*, and *C* are the integral area, number of nuclei, and concentration of the compound of interest (x) and the calibrant (cal), respectively. The concentration (g/L) of compound x can be calculated by multiplying the molecular weight of the compound.

### 2.3. Annotation of spectra

Metabolomic spectral data were normalized using the NMR reference peak such as trimethylsilyl propanoic acid (TSP) and multiple internal regions of the spectrum. (i.e., 3.41–3.47 ppm at glucose, 1.37–1.41 ppm at lactate, 1.9– 1.95 ppm at acetate, 2.36–2.39 ppm at pyruvate, 1.17–1.2 ppm at butyrate (3-HB), 1.092–1.107 ppm at valine). The sum of these internal regions has the same average value in all conditions tested. However, the spectral region corresponding to residual DI water, formate, and TSP was excluded from the metabolomic region as a reference region. Each characteristic region of the metabolite was integrated using Mnova v14.30 (Mestrelab Research, CA, USA) features. Typical 1H NMR spectra of the CAR-T cells and A549 tumor cells are shown in Fig. 1 which represent the identified metabolites by NMR spectra.

**Fig. 1.**
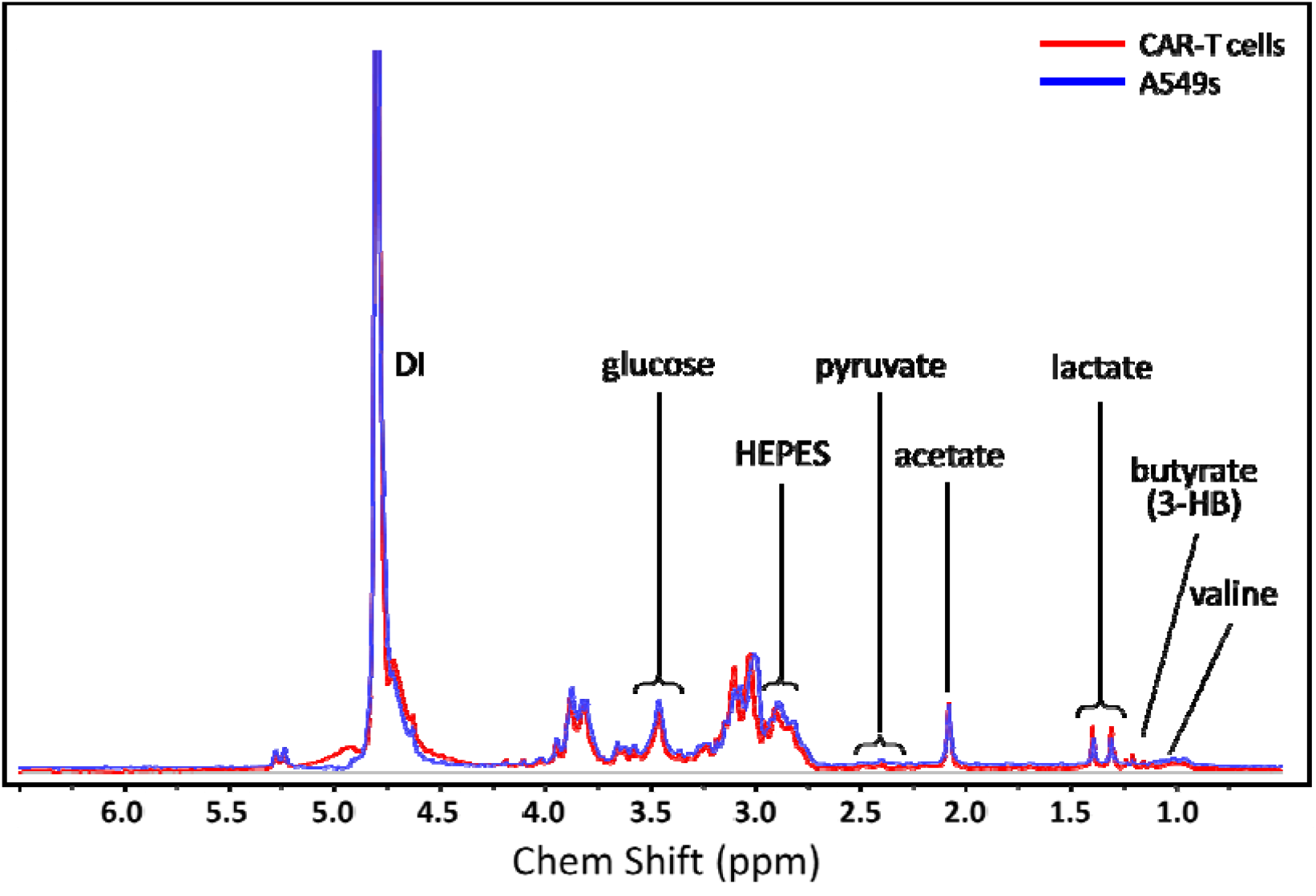
NMR spectra of typical CAR-T cells and A549 tumor cells. The spectrum indicated detective metabolites, HEPES, acetate, glucose, lactate, butyrate (3-HB), valine, and pyruvate.

### 2.4. Implementation of PCA algorithm

PCA is a dimensionality reduction method that requires a fundamental understanding of linear algebra [22]. However, the underlying concept can be explained through a geometrical interpretation of the data. Consider a matrix X with N rows and K columns, where N represents the row vector of observations and K represents the column vector of variables. The variables in the matrix correspond to the weights assigned to each original variable during the calculation of the principal component. To visualize the data, a variable space with K-dimensions is constructed (see Fig. 2(a)), where each variable represents one coordinate axis. Each variable’s length is standardized based on a scaling criterion, typically by scaling to unit variance using Z-score normalization (Gaussian). In this variable space, each observation (i.e., the Nth row in the matrix) is represented as a point in the K-dimensional space, forming a scatter of points. A variable space with K-dimensions is often employed, where each dimension represents a variable. The length of each variable axis in the space is typically standardized using a scaling criterion, such as unit variance scaling. In practice, it is common to visualize only a subset of the dimensions, such as a three-dimensional variable space. In this context, the observations in a data matrix X, which has N rows, can be represented as a scatter of points in the K-dimensional variable space.

**Fig. 2.**
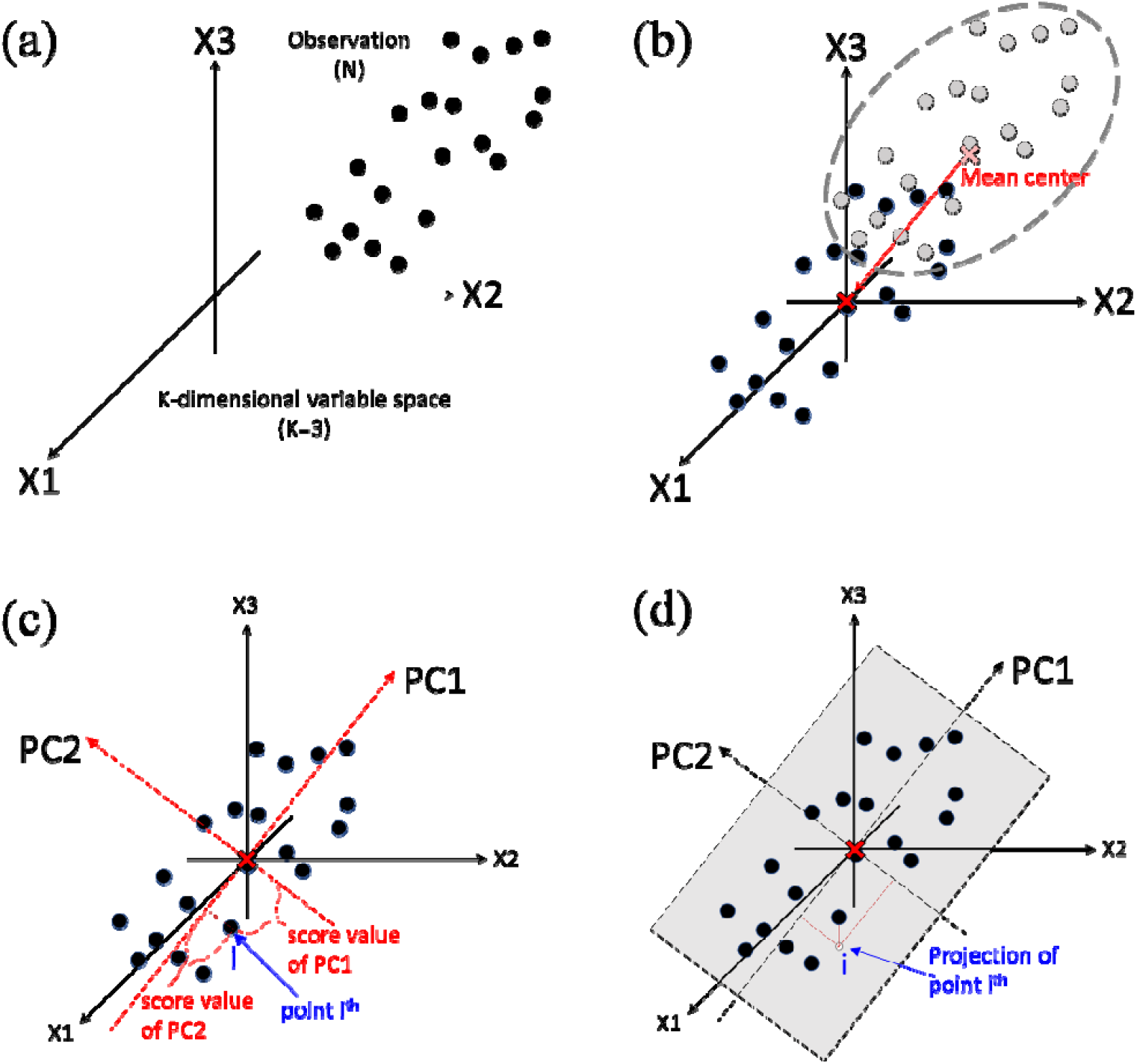
The illustration of PCA algorithm. (a) a variable space with K-dimensions is often employed, where each dimension represents a variable, (b) mean centering, (c) defined principal components, (d) two principal components define a model plane.

The process of mean-centering in data analysis entails the subtraction of the mean or average of the variables from the data in Fig. 2(b). This vector of averages represents a point in the K-dimensional variables space. The subtraction of the averages from each point of the dataset leads to a repositioning in the coordinate system in the way that the new origin corresponds to the average point. The mean-centering process involves subtracting the variables’ averages in the data from Fig. 2(a). The vector of averages can be considered as a point in the K-dimensional variables space. By subtracting the averages from the data, the coordinate system is repositioned such that the point representing the average of the variables becomes the origin. Fig. 2(c) illustrates the determination of principal components. Once all of the data has been mean-centering, and then each data was scaled to unit variance, the first principal component is computed. PC1 is the line that best accounts for the shape of the K-dimensional variable space. The line is the best approximation of the data that represents the maximum variance direction passing through the average point. Each observation is projected onto this line to obtain a coordinate value, also known as the score [23–26]. Typically, only one principal component (PC1) is insufficient to model systematic variation in a dataset, so a second principal component (PC2) is calculated. PC2 is represented by a line in the K-dimension variable space. This line is orthogonal to PC1 [27] and passes through the average point. This line improves the approximation of the data as much as possible. PC1 is the first principal component that reflects the largest amount of variance in the data. This component represents the line that best captures the shape of the points. Each data point is projected onto this line to obtain a score. While the PC2 is oriented to reflect the second largest amount of variance in the data and presented orthogonal to PC1. Like PC1, PC2 also passes through the average point of the data. Once the first two principal components have been calculated, they define a two-dimensional subspace in the K-dimensional variable space, as illustrated in Fig. 2(d). By projecting all observations onto this subspace, we can visualize the structure of datasets [28–29]. The coordinate values of the observations in this plane are called scores, and the resulting plot is known as a score plot. The two principal components obtained from PCA define a two-dimensional plane that provides a visual representation of the original multidimensional space. This projected plane allows for the visualization of the data set’s structure. The coordinates of the observations on this plane are known as scores, which can be used to plot a score plot. Finally, we implemented the PCA algorithm with the easy-to-use R package, factoextra or FactoMineR. PCA scores plot, a biplot, and box plots were generated by R-language and exported as bitmap files.

## 3. Results

### 3.1 Activities of CAR-T cells driven by distinct signaling components

We identified multiple designs that outperformed standard 4-1BB (BBz CAR) and CD28 (28z CAR) designs in a serial tumor rechallenge model for T-cell exhaustion. Based on this, to associate signaling components of CAR driving long-term persistence of CAR-T cells with metabolomic profile, we employed three different CAR ICD designs (i.e., CAR2, CAR4, and CAR10) into serial rechallenge assay where receptor tyrosine kinase-like orphan receptor 1 CAR-T (ROR1 CAR-T) cells were repeatedly challenged with A549 tumor cells endogenously expressing ROR1 over 70 days. As a proxy for assessing the long-term persistence of T cells, we measured real-time cytotoxicity, proliferation capacity, and T cell memory phenotypes (Fig. 3(a)). As a result, we observed comparable killing capacity of all three designs up to Day 20, yet CAR2 encoding CD28 (28z CAR) design started to show a decline in cell counts, whereas CAR10 encoding 4-1BB (BBz CAR) design and CAR4 encoding CD40 design maintained their proliferative capacity for a longer time. While this observation corresponds to extensively accumulated outcomes in numerous contexts displaying better persistence of BBz CAR over 28z CAR, we did not detect a significant difference in memory phenotypes of two designs where both designs exhibited an equivalent proportion of central memory (CD45RA-CD62L+) T cells (CAR2, 20.7%; CAR10, 23.5%) on Day 20 of serial rechallenge. However, we discovered that CAR4-T cells preserved a 2-fold higher proportion of central memory cells (46%) and showed consistent proliferative capacity across all replicates up to Day 70 (Fig. 3 (b) and (c)). We reasoned that the metabolomic profile of relatively more persistent CAR4 would be different from those of CAR2 and CAR10. To discover a metabolomic signature of CAR4, we collected supernatant of tumor cell co-culture on Day 20 and profiled them via benchtop NMR.

**Fig. 3.**
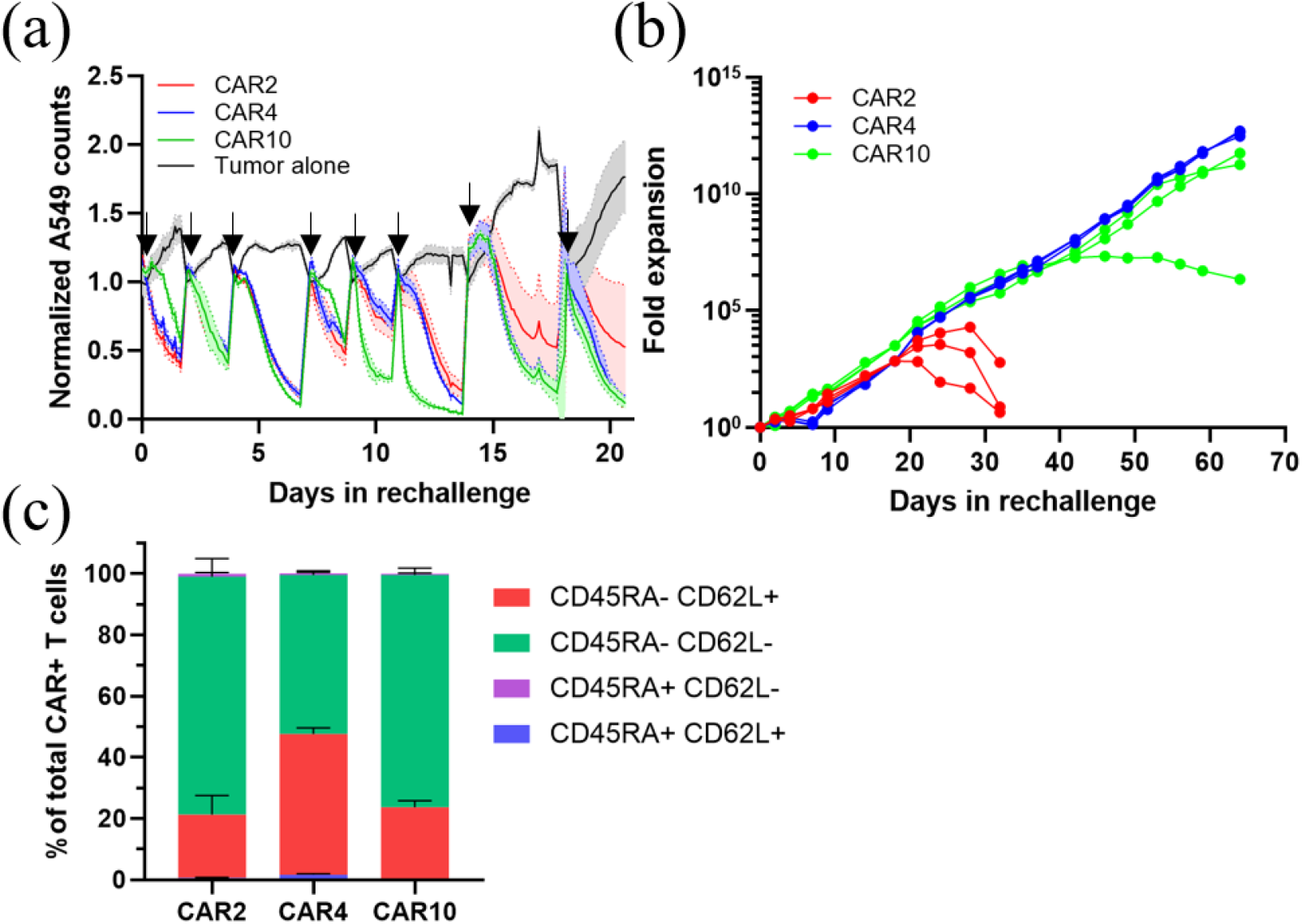
Different signaling components of CAR lead to diverse functional outcomes in T cells. (a) Plot describing repetitive cytotoxic features of CAR-T cells during the serial rechallenge assay with A549 tumor cells. Arrows indicate time points of tumor challenge. (b) Graph showing fold expansion of CAR-T cells over 70 days of repeated tumor challenge. Each line represents each replicate. (c) Bar graph depicting memory phenotypes of CAR-T cells on Day 20 of serial rechallenge assay. (where n = 3 and error bars represent standard deviation)

### 3.2 Quantitative variation of metabolites among CAR-T cells

Fig. 4 shows a clear quantitative variation in the metabolite profiles between the cancer cells and CAR-T cells. Glucose was significantly higher (p <0.1) in A549 tumor cells, that is because cancer cells have high metabolic demands. Clinically, cancer cell metabolism is characterized by enhanced glucose uptake and utilization. Lactate was significantly higher (p <0.1) in CAR4s, compared to CAR2s, and CAR10s. Butyrate, valine, and pyruvate were found to be more abundant in the CAR2s and CAR4s but not in the other CAR10s. CAR10s were significantly higher (p <0.1) in acetate, compared to CAR2s and CAR4s.

**Fig. 4.**
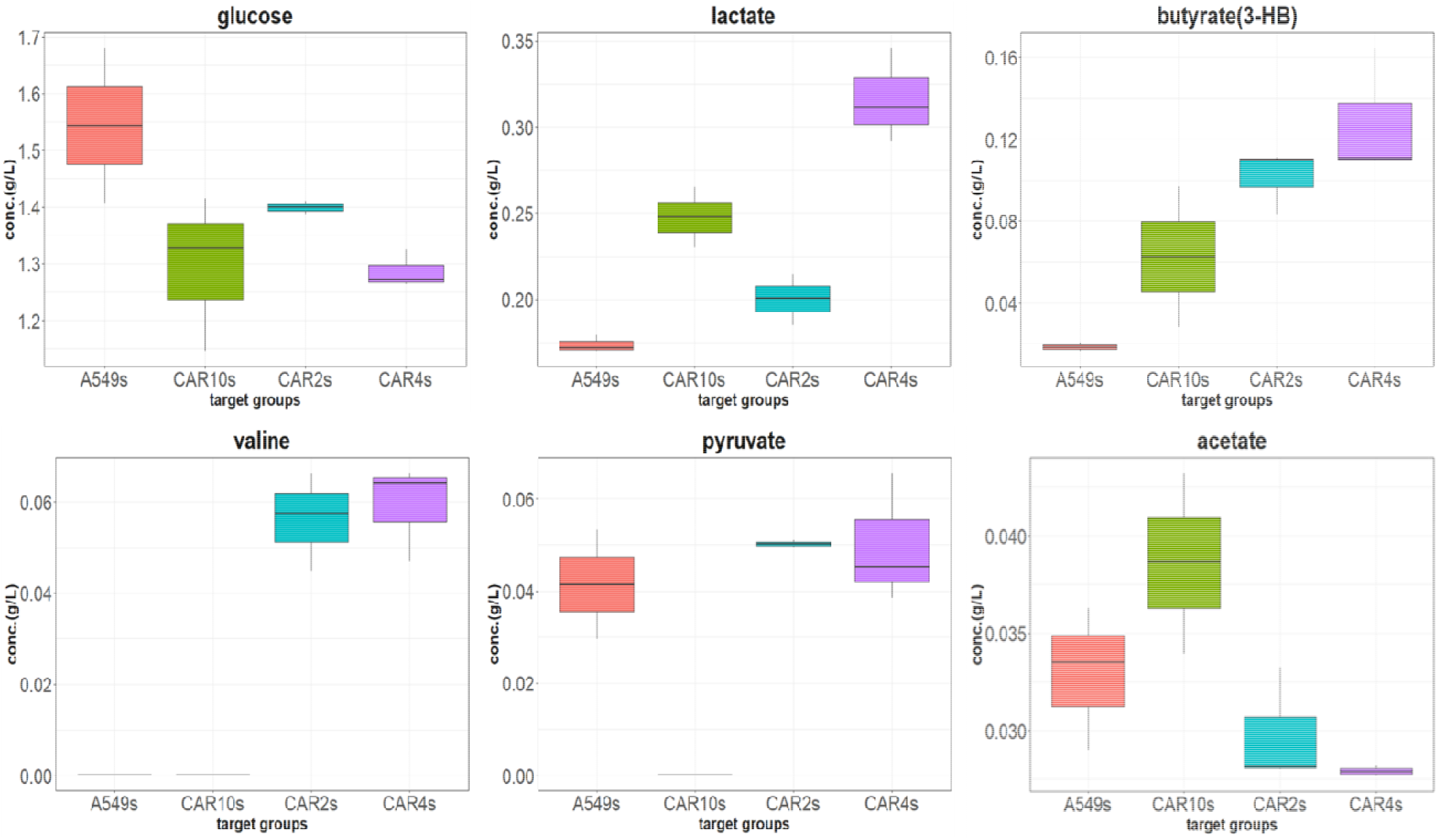
Quantitative variation of metabolites among CAR-T cells

### 3.3 Principal components analysis of CAR-T and tumor cell metabolites

The model has six variables (i.e., metabolites: acetate, glucose, lactate, butyrate(3-HB), valine, pyruvate) and twelve targets (i.e., cells: A549-1, A549-2, A549-3, CAR10-1, CAR10-2, CAR10-3, CAR2-1, CAR2-2, CAR2-3, CAR4-1, CAR4-2, CAR4-3). HEPES was detected but not included in metabolites, because it is commonly used in cell culture media as a buffering substance. The principal component analysis was performed to reduce variables by extracting principal components from the data. In principal component analysis, the eigenvalues measure the amount of variation retained by each principal component. We examine the eigenvalues to determine the number of principal components to be considered. The eigenvalues can be used to determine the number of principal components, whereas variances can provide all the information of principal components. Fig. 5 is the Scree plot showing the acceptable value of the amount of data variation according to dimensionality reduction by extracting principal components of data. In our analysis, the correlation between the metabolites and the targets was set as two-dimensional space (PC1 and PC2). The first two principal components explain 85.1% (= PC1(53.5%) + PC2(31.6%)) of the variation. This is an acceptable percentage

**Fig. 5.**
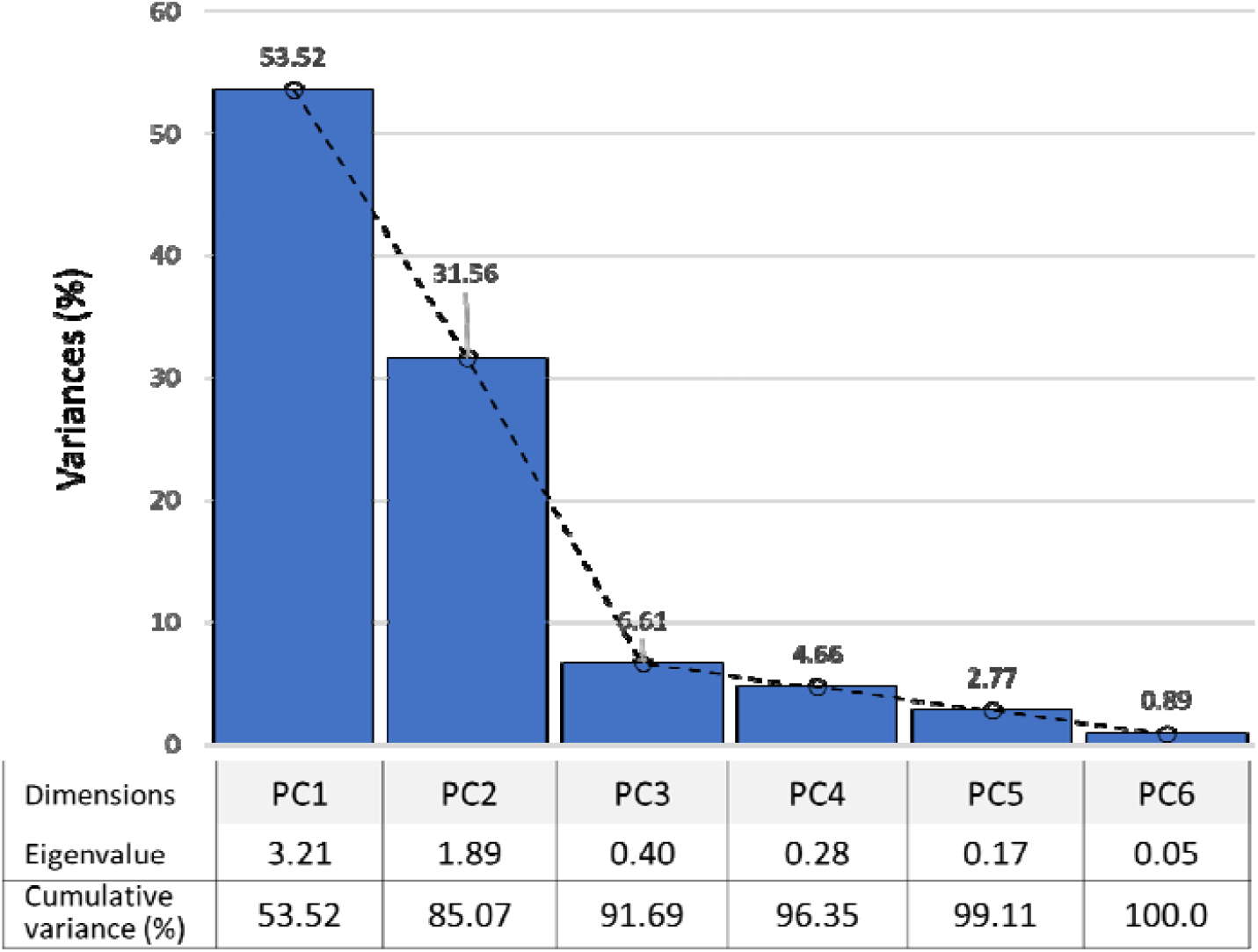
The scree plot shows the ratio of data variation to dimensionality reduction of the model.

### 3.4 Main contributions of the metabolites to the PC1 and PC2

The contributions of metabolites in accounting for the variability in a given principal component are expressed in a two-dimensional plot. Metabolites that are correlated with PC1 and PC2 are the most important in explaining the variability in the data set. The other metabolites that do not correlate with any PC or correlate with the last dimensions are variables with low contribution and might be removed to simplify the overall analysis. The total contribution of a given metabolite, on explaining the variations retained by two principal components, is calculated as contribution (%) = [(C1 * Eig1) + (C2 * Eig2)]/(Eig1 + Eig2), where C1 and C2 are the contributions of the variable on PC1 and PC2, respectively and Eig1 and Eig2 are the eigenvalues of PC1 and PC2, respectively. The expected average contribution of a variable for PC1 and PC2 is [(6* Eig1) + (6 * Eig2)]/(Eig1 + Eig2) = [(6*3.21)+(6*1.89)]/(3.21+1.89) = 6 %. The red dashed line indicates the expected average contribution in Fig. 6 (a) and (b). If the contribution of the metabolites were uniform, the expected value would be 1/length(metabolites) = 1/6 = 16.7%. For a given component, a metabolite with a contribution larger than this cutoff could be considered as important in contributing to the component. Herein, it can be seen that the metabolites (i.e., valine, pyruvate, and butyrate) contribute the most to the PC 1 and PC2

**Fig. 6.**
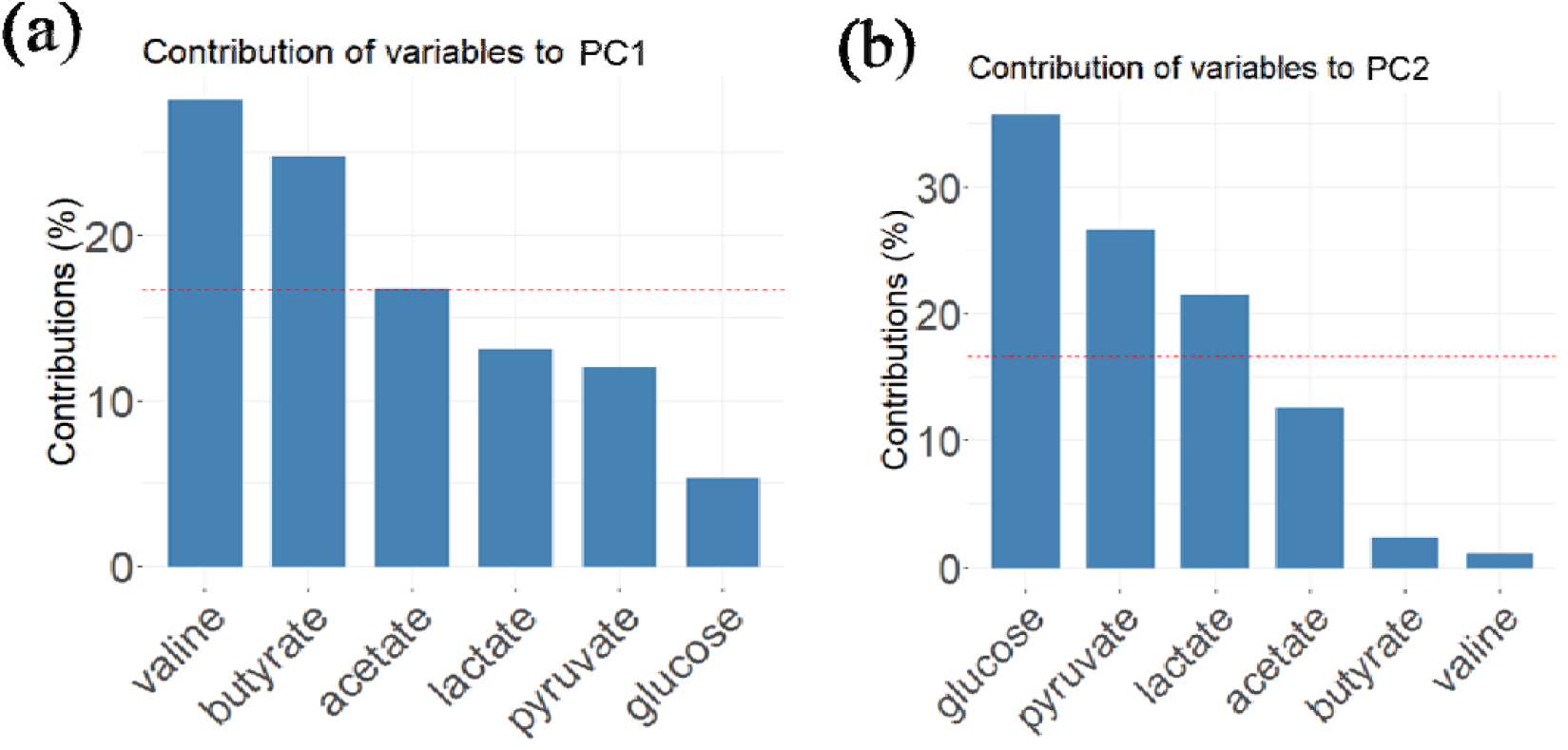
Contribution of metabolites to the principal component, (a) PC1-axis and (b) PC2-axis.

### 3.5 Correlations similarity among metabolites

The contribution of among six metabolites on the component axes, Fig. 7 (a) and (b) are used to assess the quality of the representation of the metabolites of the principal component. The larger the value of the contribution, the more the metabolite contributes to the component. To find out the correlations among metabolites from CAR-T cells and A549 tumor cells, they were clustered by classifying the metabolites into four groups using the k-means clustering algorithm. The K-means clustering was analyzed for similarity by measuring the distance between metabolites of the plot. As a result of categorization through the K-means clustering algorithm that tries to group a correlated strength of CAR-T cell metabolites in the form of clusters, four clusters were formed such as butyrate, lactate is cluster 1, glucose is cluster 2, acetate is cluster 3, and pyruvate, valine is cluster 4, respectively. Metabolites within each cluster are positively correlated, when the value of one metabolite increases or decreases, the value of the other variable has a tendency to change, while cluster 1 and cluster 2 are inversely correlated, meaning that when the metabolism of cluster 1 works in opposite ways for cluster 2 and vice versa. Also, cluster 3 and cluster 4 are the same way.

**Fig. 7.**
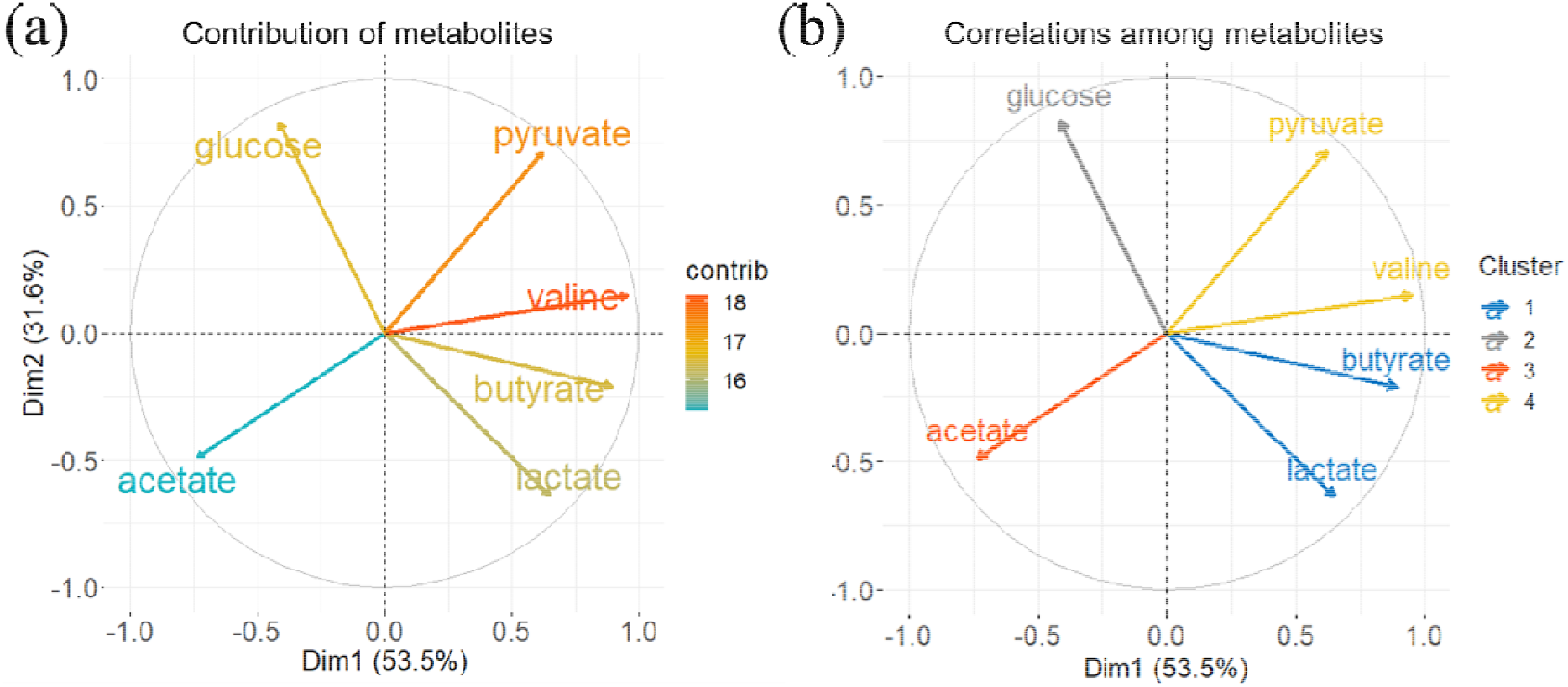
Correlations similarity among metabolites. (a) contribution of among six metabolites, (b) four different clusters between CAR-T cell metabolites. (where the contrib’s index represents the contribution of metabolites to the principal component)

### 3.6 Profiling of CAR-T and tumor cell metabolites

The profiling of metabolites aims to interpret the relationship between metabolites and target cells. Fig. 8 presents the relationships among metabolites and target cells including nine CARs and three A549 tumor cells. There were distinct metabolic differences among target cells such as CAR2s, CAR4s, and CAR10s groups. The distance of a metabolite from the plot origin represents its impact on the model. Pyruvate and valine are located far from the origin, indicating that they have a strong impact on the model. On the other hand, the locations of targets on the principal components plane reveal their similarity in terms of metabolite profiles. Targets that are close to each other are correlated, while those far apart are not. From this plot, metabolites with similar information are grouped based on their correlations. A clearer insight into distinctly different states of CARs on the metabolite profile was gained via contribution values to the principal components. The PCA plot of CAR-T and tumor cell metabolites correlation demonstrated clear variation in the metabolite profiles among the CARs groups and the A549 group. CAR4 group was inversely correlated A549 group on the PCA plot. Glucose was high in the A549 group, while lactate was significantly higher in the CAR4 group. In other words, since A549 has a strong tendency to change metabolism by increasing glucose uptake and fermenting glucose into lactate, CAR4 cells with enhanced glycolysis can be considered to be able to inhibit cancer cell growth. The CAR2 group was inversely correlated CAR10 group on the PCA plot, too. Acetate was found to be higher in the CAR10 group, while pyruvate was higher in the CAR2 group. In previous studies, 28z CAR (CAR2) and BBz CAR (CAR10) showed significantly different metabolic profiles, and the amount of pyruvate that was from a metabolite of glycolysis of CAR2 was significantly higher than that of CAR10.

**Fig. 8.**
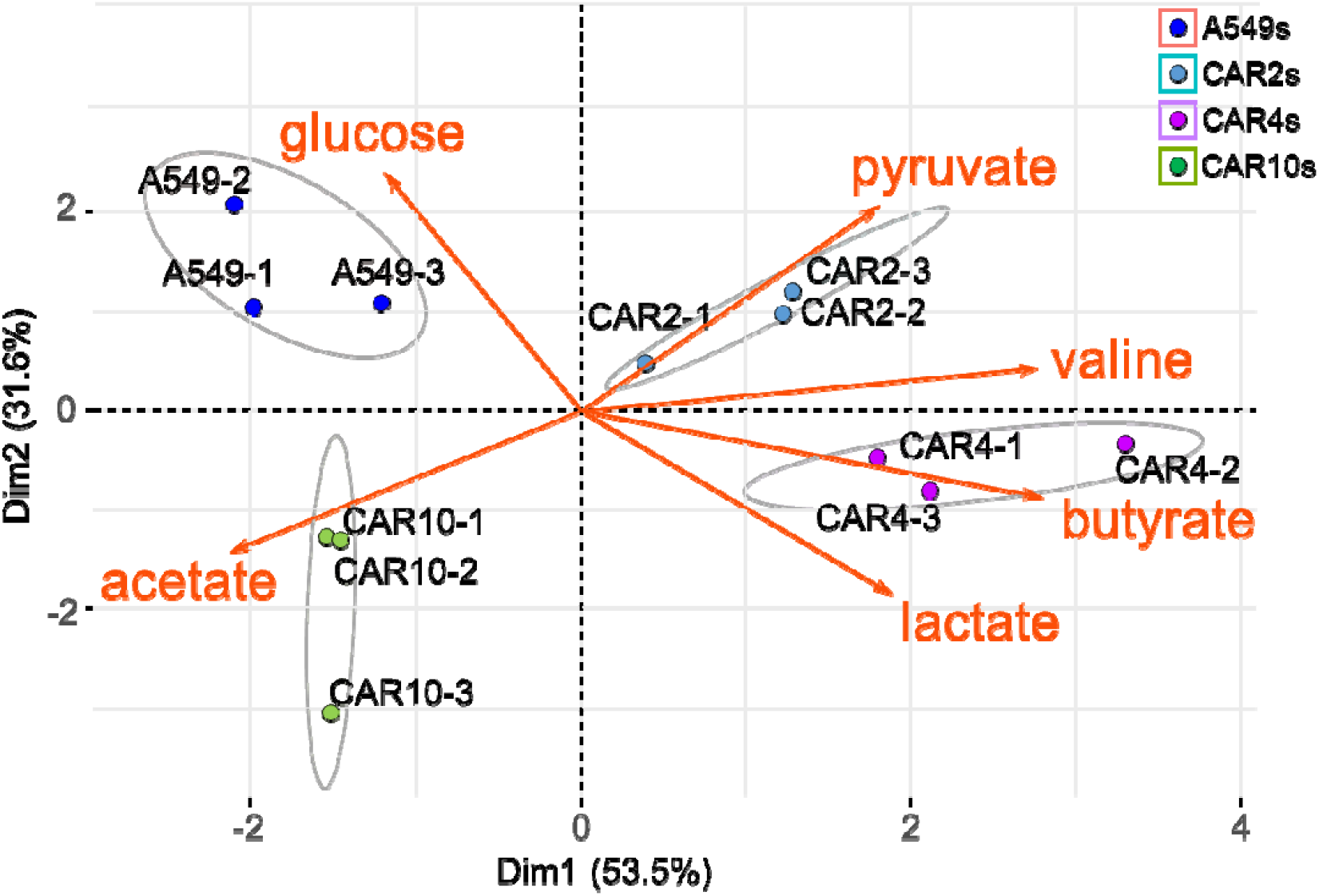
Profiling of CAR-T and tumor cell metabolites to the principal components. There is shown in the relationship between metabolites and target cells according to metabolite contribution values to influential target cells. (where group fill colors represent the correlated target cells, ellipse: confidence = 0.99)

## 4. Discussion

PCA was used to dissect multi-variable datasets generated by a benchtop NMR instrument. The simple workflow outlined in this article offers a promising avenue for investigating the metabolites of CAR-T cells, with broader applicability to other biological cells as well. To make sense of such datasets, particularly diverse NMR spectra data sets from different types of CAR-T cells, PCA proved indispensable. It efficiently reduces data dimensionality, making it more interpretable while retaining essential information. The profiling plot (see Fig. 8) transforms the NMR spectra data’s principal components domain into actionable metabolic insights. This transformation serves as a crucial tool for deciphering intricate biological data.

In this study, we harnessed PCA to minimize the impact of extraneous features and enhance model interpretability for CAR-T cells and their metabolites. Notably, PC1 and PC2, as depicted in Fig. 6 (a) and (b), collectively accounted for 85.1% of the variance. It’s worth noting that proper data scaling is critical for PCA’s accuracy, particularly in two or three dimensions. PCA findings unveil significant disparities in the metabolic profiles of CAR-T cells contingent on distinct CAR ICD designs. For instance, CAR4-T cells exhibited a remarkable 2-fold higher proportion of central memory cells and consistently maintained proliferative capacity for over three times longer, as demonstrated over 70 days in Fig. 3. In other words, CAR4-T cell metabolic specificity (CD40 signaling) against A549 tumor cells appears to have resulted in the consistent proliferative ability of CAR4 by creating differences in metabolic pathways and expression levels of major genes. This suggests that the metabolic profile of CAR4 could potentially serve as a robust biomarker for CAR-T cells with robust immunity in future developments. Furthermore, our PCA investigations provide compelling evidence that newly engineered CAR signaling domains have a discernible impact on T cell behavior and metabolic reprogramming, underscoring the concept of metabolic plasticity. In various analytical and omics disciplines, multi-dimensional visualizations using principal components are standard practice, either by projecting samples onto the components or by plotting them based on their correlation with these components. Nevertheless, it’s important to acknowledge that reducing dimensionality can lead to information loss, necessitating experimentation with different combinations of components when visualizing a dataset. As the growth of biological big data from diverse sources only accelerates, PCA is poised to play a pivotal role in uncovering new applications in the future.

## Author contributions

S.H.S. and H.R. designed the outline of an article and drafted the manuscript. S.H.S. analyzed the metabolome NMR data using PCA. J.H. set up the NMR spectroscopic system and measured metabolites from the CAR-T cell samples. T.K. designed the CARs and edited the manuscript in the parts of CAR design sections. J.M., J.R., and S.E. designed the study and prepared CAR-T cell samples. All authors read and approved the manuscript.

